# ANP rescues the preeclamptic symptoms in *Axl* knockout mice by improving trophoblast function ANP rescues the preeclamptic symptoms

**DOI:** 10.1101/2023.04.14.536838

**Authors:** Chan Zhou, Yunqing Zhu, Liang Zhang, Xuexiang Li, Jiatong Yao, Miaomiao Zhao, Zi-jiang Chen, Cong Zhang

**Affiliations:** Center for Reproductive Medicine, Ren Ji Hospital, School of Medicine, Shanghai Jiao Tong University, Shanghai,200135, China; Shanghai Key Laboratory for Assisted Reproduction and Reproductive Genetics, Shanghai, China; Shandong Provincial Key Laboratory of Animal Resistance Biology, College of Life Sciences, Shandong Normal University, Jinan, Shandong, 250014, China; Research Center of Translational Medicine, Jinan Central Hospital Affiliated to Shandong First Medical University, Jinan, Shandong 250013, China; Center for Reproductive Medicine, Cheeloo College of Medicine, Shandong University, Jinan, Shandong, 250012, China; National Research Center for Assisted Reproductive Technology and Reproductive Genetics, Shandong University, Jinan, Shandong, 250012, China

**Keywords:** ANP, AXL, CORIN, decidua, placenta, preeclampsia

## Abstract

Pre-eclampsia (PE) is characterized by maternal hypertension and/or proteinuria after 20 weeks of gestation. Shallow trophoblast invasion into spiral artery (SpA) and defective decidualization have been implicated in the pathogenesis of PE. However, the underlying mechanisms remain unclear. AXL receptor tyrosine kinase is a biomarker and therapeutic target for a variety of metastatic cancers while tumorigenesis and placentation share many features. In this study, a new function of AXL in promoting SpA remodeling and trophoblast invasion were demonstrated. The pregnant *Axl* knockout (*Axl*^−/−^) mice exhibited typical PE symptoms, including increased blood pressure and proteinuria. Artificially induced decidualization experiments showed that appeared normal, however, RNA-seq results of *Axl*^−/−^ deciduoma revealed abnormal expression of a number of transcripts, including *Corin*, which encodes a cardiac protease that activates atrial natriuretic peptide (ANP). Treatment with ANP reversed the PE symptoms. Moreover, in decidua from women afflicted with PE, AXL level was significantly lower than that in normal pregnancies. These data show that the abnormality of decidua-derived AXL-CORIN-ANP affects maternal-fetal crosstalk and contributes to PE.

## Introduction

Pre-eclampsia (PE) is a gestational syndrome affecting 5-8 % of all pregnancies, and is the leading cause of fetal and maternal morbidity and mortality. PE is characterized by the newly occurred hypertension and/or proteinuria after 20 weeks of gestation (Rana, Lemoine et al., 2019, Souza, Gülmezoglu et al., 2013). The only current treatment is the delivery of the placenta (Burton, Redman et al., 2019). The etiology of PE is heterogeneous and complex, and studies on PE in humans have hitherto identified endothelial dysfunction, immune dysregulation and defective decidualization as elements in the pathogenesis of this disease (Garrido-Gomez, Quiñonero et al., 2020, Ho, van Dijk et al., 2017, Lv, Tong et al., 2018, Rong, Yan et al., 2021, Zhou, Zhang et al., 2008). Animal models, particularly mice and rats, have been used to study the molecular mechanisms underlying PE (Blois, Dechend et al., 2015, Winship, Koga et al., 2015).

Shallow trophoblast invasion of the uterine spiral artery (SpA) leads to abnormal placentation followed by the maternal symptoms in clinical manifestation (Jim and Karumanchi, 2017). It is widely believed that PE appears in 2-stage pathogenesis in that the shallow trophoblast invasion of the uterine decidua leading to abnormal placentation followed by the maternal symptoms (Fisher, 2015). The insufficient arterial modification by trophoblasts results in poor perfusion of the placenta with maternal blood, leading to PE and/or intrauterine growth restriction of the fetus (Siddiqui, Nandi et al., 2016). Decidua formation precedes conceptus implantation and decidualized cells are a source of secreted molecules such as growth factors, cytokines, metalloproteases, and protease inhibitors that support/control trophoblast invasion (Garrido-Gomez, Dominguez et al., 2011, Jovanović & Vićovac, 2009).

AXL is a receptor tyrosine kinase belonging to the Tyro3-Axl-Mertk family, with GAS6 is the common ligand. The binding of GAS6 to AXL stimulates AXL dimerization and intracellular domain autophosphorylation, thereby transducing extracellular signals to downstream effectors, playing essential roles in cell adhesion, invasion, migration, proliferation, and pro-inflammatory cytokine production (Graham, DeRyckere et al., 2014, Linger, Keating et al., 2008). AXL is expressed in human ovarian tumors, and the inhibition of AXL signaling results in decreased cell invasion and prevents ovarian tumor progression (Rankin, Fuh et al., 2010). AXL has been described as a biomarker and therapeutic target for a variety of metastatic cancers (Holland, Powell et al., 2005). Despite the known role for AXL in tumor cell invasion, its relationship with pregnancy and PE remains unknown.

In the present study, we investigated the potential physiological role(s) of AXL during pregnancy, by examining of *Axl* knockout (*Axl^−/−^*) mice. We found that *Axl* knockout caused PE symptoms in pregnant mice bearing this mutation. *Axl* knockout affected the interaction between maternal decidual tissue and placental trophoblast, causing compromised trophoblast invasion and SpA remodeling. This leads to increased blood pressure, proteinuria, abnormal renal glomerular structures and reduced fetal weight. Cross mating and embryo transplantation confirmed maternal uterine tissue is the origin of this PE symptoms. RNA-seq results demonstrated the significantly decreased *Corin* expression in *Axl*^−/−^ decidua. CORIN is a cardiac protease that activates atrial natriuretic peptide (ANP), a cardiac hormone that regulates blood pressure and sodium homeostasis. ChIP-qPCR demonstrated that decreased CORIN in *Axl*^−/−^ decidua resulted from reduced STAT3 binding to the *Corin* promoter, and therapeutic intervention with ANP could rescue the PE symptoms of *Axl^−/−^* pregnant mice. Moreover, we found that AXL levels were decreased in the decidua of preeclamptic women compared to those in normal pregnancies. Thus, our study reveals that AXL-related decidual dysfunction impairs the maternal-fetal crosstalk, and is a factor in PE pathogenesis.

## Results

### *Axl* knockout leads to PE symptoms in pregnant mice

To assess the potential impact of AXL during pregnancy, we examined a previously established mouse model that employed homologous recombination to delete exon 9 of the *Axl* locus (Lu, Gore et al., 1999). We found that the pregnant *Axl^−/−^*mice had significantly increased systolic blood pressure compared to pregnant *Axl^+/+^* mice, beginning approximately embryonic day 14.5 (E14.5), and blood pressure continued to rise as pregnancy ensued and delivery approached (Figure 1A). This elevated pressure resembles the early onset gestational hypertension observed in preeclamptic women (Rana et al., 2019). Elevated urinary albumin/creatinine ratios are a key symptom of PE, and urine collected from E14.5 to E17.5 for 72 h in metabolic cages indicates that the pregnant *Axl^−/−^* females had significantly elevated urinary albumin/creatinine ratios compared to the pregnant *Axl^+/+^* mice (Figure 1B).

**Figure 1.**
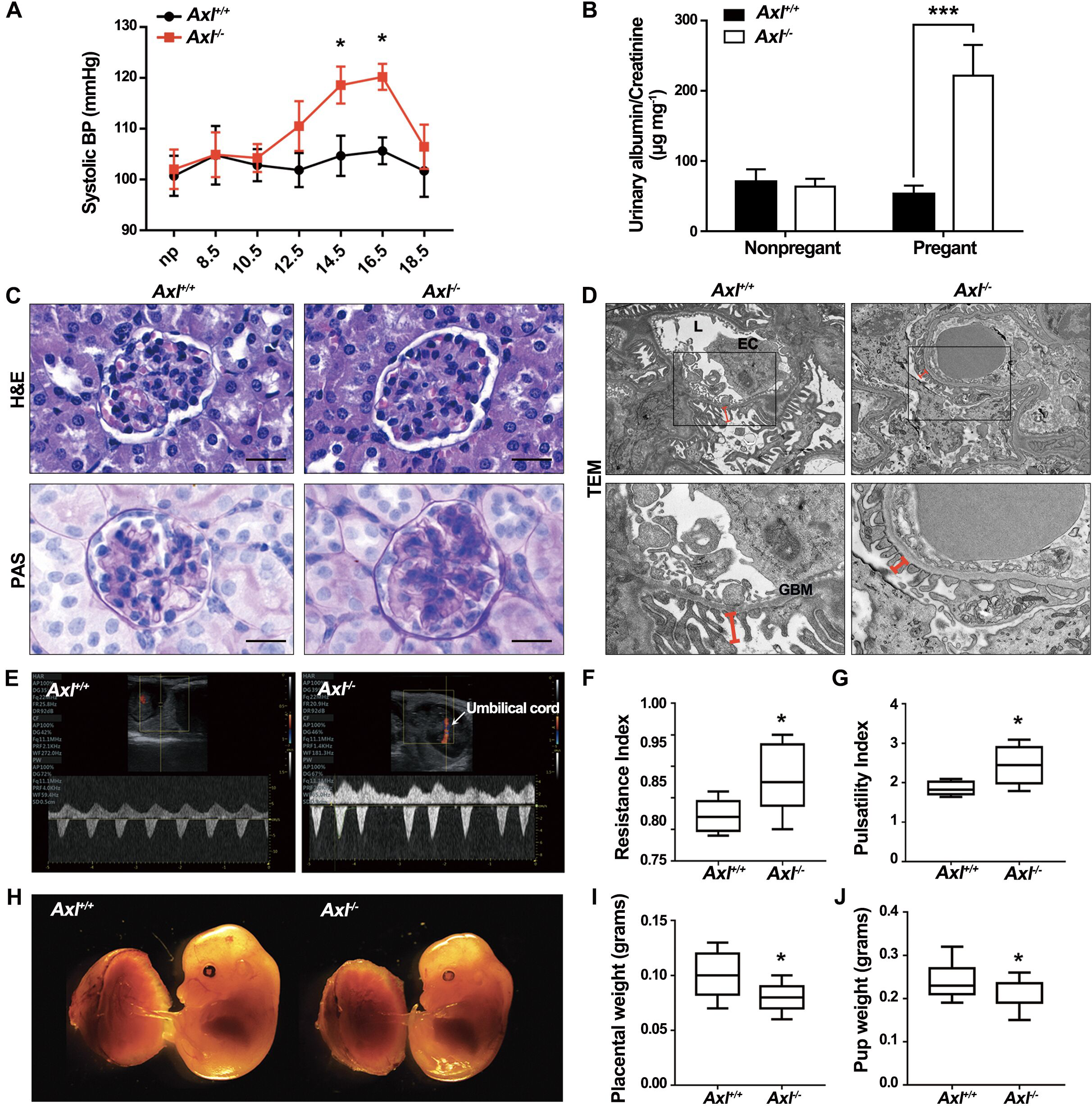
Hypertension, proteinuria, renal pathology, and reduced fetal weight in pregnant *Axl* knockout mice. (A) Repeated tail-cuff systolic blood pressure (BP) measurements of *Axl^+/+^*and *Axl^−/−^* mice before and during pregnancy. (B) Urine albumin/creatinine ratios of *Axl^+/+^* and *Axl^−/−^* mice, urine samples were collected (from E14.5 to E17.5) in metabolic cages. (C) H&E and PAS staining of renal tissues from pregnant *Axl^+/+^* and *Axl^−/−^* mice. The H&E staining reveals glomerular enlargement in *Axl^−/−^* mice. The PAS staining shows increased extracellular matrixes and collapsed glomerular capillaries in the *Axl^−/−^*mice. (D) Transmission electron microscopy (TEM) imaging of the narrow glomerular capillary lumen (L), thick glomerular basement membranes (GBM), and the shortened secondary processes of podocytes indicates endothelial hyperplasia and extracellular matrix expansion in *Axl^−/−^*glomeruli (indicated by red lines). X-magnification 15,000x and Y-magnification 6,000x. (E) Uterine artery blood flow velocity waveforms in *Axl^+/+^*and *Axl^−/−^* mice, assessed using pulse Doppler ultrasonography starting from E14.5. (F and G) Uterine artery waveforms from *Axl^+/+^*and *Axl^−/−^* were analyzed to obtain resistance and pulsatility indexes. Values are means ± SE. Data are from 4–5 independent litters. (H) Representative photographs of placentas and fetuses of E14.5 *Axl^+/+^* and *Axl^−/−^* mice. (I-J) Placental and fetal weights (E14.5) were reduced in *Axl^−/−^* females compared to *Axl^+/+^* females. All data are expressed as means ± SD). Scale bar, 20 μm. np, not pregnant. *, *p* < 0.05; **, *p* < 0.01; ***, *p* < 0.001.

Analysis of kidney glomerular sections taken at E14.5 revealed that the *Axl^−/−^* mice had ischemic glomeruli, as evidenced by reduced numbers of red blood and cells in the enlarged glomeruli (Figure 1C). Further, transmission electron microscopy of kidney glomerular sections revealed signs of glomerular hyperplasia in the pregnant *Axl^−/−^*mice, mainly manifested as capillary lumen narrowing, capillary loop occlusion, endothelial cell swelling and hypertrophy, and shortening of podocyte secondary processes (Figure 1D). These histopathology features recapitulate the renal pathology of women suffering from PE (Mol, Roberts et al., 2016, Strevens, Wide-Swensson et al., 2003).

We also conducted non-invasive Doppler ultrasound of the umbilical artery to assess the intrauterine blood supply (monitoring umbilical blood flow and placental resistance). Pulse wave images were acquired from the spiral artery in the Doppler mode and used to quantify the velocity of umbilical artery blood flow. *Axl^−/−^*females had significantly elevated uterine artery resistance index values and significantly elevated uteroplacental pulsatility index values compared to *Axl^+/+^* females at E14.5 (Figure 1E-1G). Additionally, pregnant *Axl^−/−^* mice had significantly reduced placental weight and fetal weight at E14.5 (Figure 1H-1J). Together, these observations indicate that pregnant *Axl^−/−^*mice developed PE symptoms.

### Impaired trophoblast invasion, SpA remodeling, and glycogen deposition in *Axl^−/−^*mice

During pregnancy, trophoblast invasion and uterine SpA remodeling have been shown to lower maternal vascular resistance and increase uteroplacental blood flow. Impaired trophoblast invasion and uterine SpA remodeling are associated with PE symptoms (Ridder, Giorgione et al., 2019). Cytokeratin staining of the uteroplacental interface showed that trophoblasts invaded deep in the decidua of *Axl^+/+^* mice at E10.5 (arrows) and that *Axl^+/+^*uterus had enlarged SpAs surrounded by trophoblasts at E14.5 (solid box) (Figure 2A, a, c). In contrast, trophoblast invasion for *Axl^−/−^* uterus was markedly reduced and only smaller arteries (solid box) existed in both superficial and deep decidua (Figure 2A, b, d). At E18.5, the *Axl^+/+^* mice displayed significantly more trophoblasts, in the decidua and in the myometrium compared to *Axl^−/−^* mice (Figure 2B, a, b). Immunofluorescence of smooth muscle actin showed that the expected replacement of smooth muscle by invasive trophoblasts in the *Axl^+/+^* deep decidua; this was not evident in the *Axl^−/−^* sections (Figure 2B, c, d). We conclude that *Axl* knockout impairs trophoblast invasion and uterine SpA remodeling in mouse maternal decidua.

**Figure 2.**
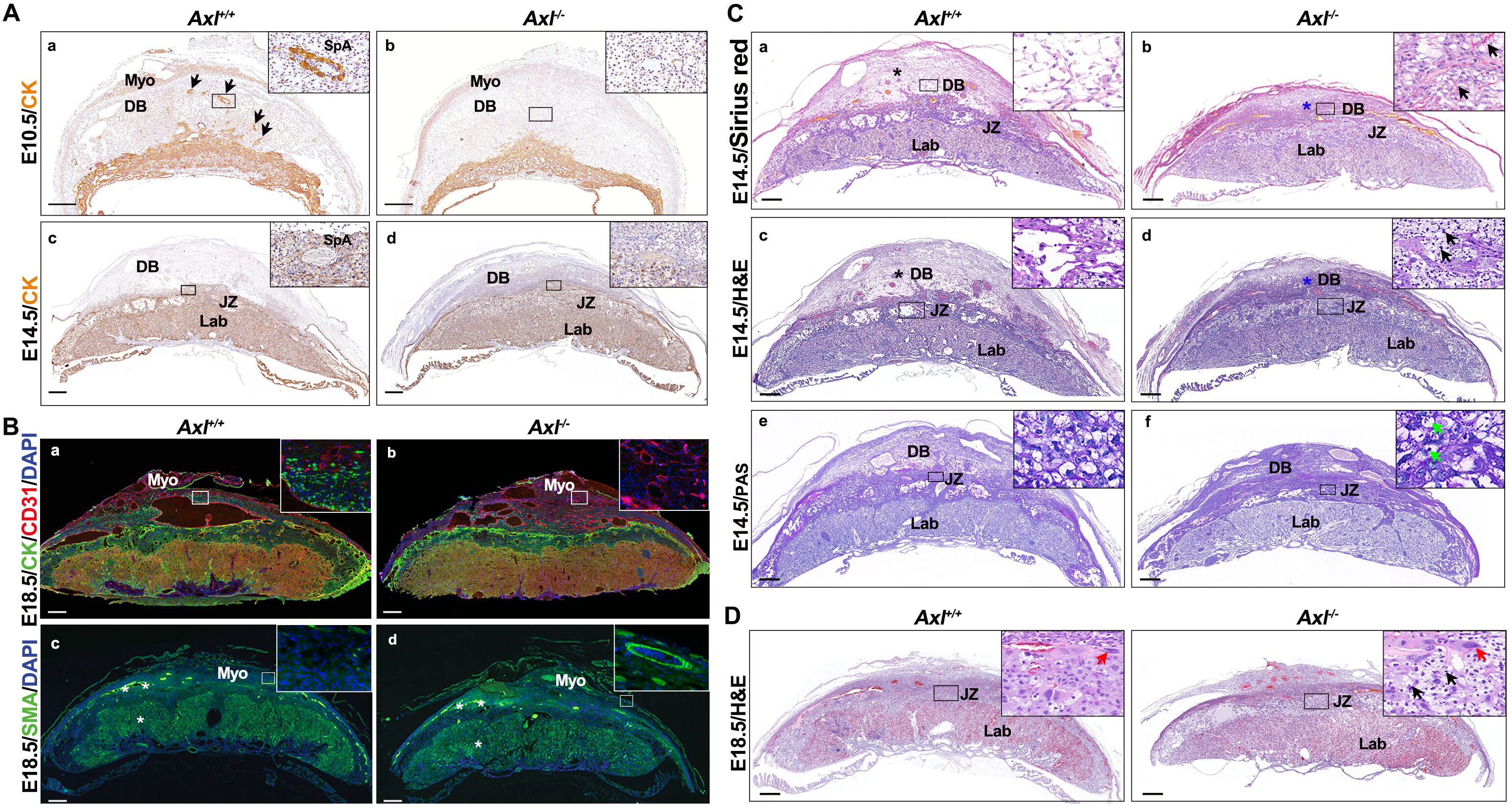
Utero-placental interfaces of *Axl^+/+^* and *Axl^−/−^* mice. (A) Immunohistochemical analysis of the *Axl^+/+^* and *Axl^−/−^* interface at E10.5 and E14.5 shows the invasion level of cytokeratin-expressing trophoblast cells (brown signal) (a-d). (B) Expression of trophoblast cell marker (cytokeratin, CK), blood vessel endothelial marker (CD31), and smooth muscle marker (smooth muscle actin, SMA), at E18.5 maternal-fetal interface of *Axl^+/+^* and *Axl^−/−^* mice. (C) Comparison of Sirius red, H&E and PAS staining of *Axl^+/+^* and *Axl^−/−^* interfaces shows a compact decidua (blue asterisks) and glycogen deposition in junctional zone (arrows) in *Axl^−/−^* mice at E14.5 (a-f). Arrows points to foci of glycogen deposition. (D) Comparison of H&E staining of *Axl^+/+^* and *Axl^−/−^* interfaces shows that glycogen deposition at E18.5 in *Axl^−/−^* mice. Red arrows indicate trophoblast cells of junctional zone. Black arrows point to foci of glycogen deposition. Scale bars, 0.5 mm. Myo, myometrium; DB, decidual basalis; JZ, junctional zone; SpA, spiral artery; Lab, labyrinth.

We also noted that the *Axl^+/+^* decidua basalis was loose (black asterisk) with reduced extracellular matrix content (Figure 2C, a, c), while that of *Axl^−/−^* was compact (blue asterisk) which contain high content of extracellular matrix (marked by arrows) (Figure 2C, b, d) by Sirius red staining of E14.5 sections. Moreover, the H&E staining of *Axl^−/−^* mice showed an apparent increase in glycogen deposition (marked by arrows) in the junctional zone (Figure 2C, d), which resembles the glycogen deposition reported for preeclamptic placentas in both human studies and animal models (Akison, Nitert et al., 2017, Cubro, Nath et al., 2021). PAS staining confirmed that glycogen deposition was present in the junctional zone of *Axl^−/−^* placentas (marked by green arrows) but not in the *Axl^+/+^* samples at E14.5 (Figure 2C, e, f). The glycogen deposition (black arrows) was still observed in the junctional zone of E18.5 *Axl^−/−^* females (Figure 2D). These data demonstrate profoundly abnormal decidua basalis structure and increased glycogen deposition in *Axl^−/−^* mice.

To further investigate whether the abnormality of the placenta was confined to the maternal-fetal interface, we conducted dual fluorescence analysis of CD31 and cytokeratin to mark blood vessels and trophoblasts in the placental labyrinth (Figure S1A, B), no differences were detected between the *Axl^−/−^* and *Axl^+/+^* samples. We conclude that insufficient trophoblast invasion, spiral artery remodeling, and aberrant glycogen deposition are restricted in decidua and the junctional zone of the placenta and are responsible for the abnormal maternal-fetal interface observed in the *Axl*^−/−^ mice.

### Maternal *Axl* knockout causes the PE symptoms due to defective decidualization in *Axl^−/−^* mice

Whether PE is paternally or maternally originated has long been debated, due to mutations that are maternally active, paternally imprinted genes normally expressed in placentogenesis (Graves, 1998). We therefore performed a variety of crosses with paternal or maternal mutations, seeking to identify the maternal-fetal interactions and the contributions of each part to the PE symptoms. *Axl^+/-^* or *Axl^−/−^*females were crossed with *Axl^+/+^* or *Axl^−/−^* males, thus generating interfaces representing four distinct genotypes. H&E staining showed that, regardless of the embryonic genotypes, the *Axl^+/-^*maternal decidua (black asterisks), displayed the normal incompact phenotype and containing enlarged SpAs (Figure 3A, a, b). In contrast, *Axl^−/−^* decidua (carrying either *Axl^+/-^* or *Axl^−/−^* fetus), were abnormal, presenting narrow vessels and compact decidual cells and a non-degraded extracellular matrix (blue asterisks). The junctional zone of *Axl^−/−^* placentas displayed excessive glycogen deposition (arrows) (Figure 3A, c, d). We also adopted an embryo transfer strategy to transfer *Axl^+/+^* embryos into *Axl^−/−^*uteri, and *vice versa* (Figure S2A). H&E staining showed glycogen deposition and a thinned junctional zone in the *Axl^−/−^* mice carrying *Axl^+/+^* fetus; these symptoms were not observed in *Axl^+/+^* mice carrying *Axl^−/−^* fetus (Figure S2B). The glomerular structures of the pregnant *Axl^−/−^* mice carrying *Axl^+/+^* fetus exhibited the same phenotypes as those of *Axl^−/−^* mice carrying the *Axl^−/−^* fetus (Figure S2C). These data demonstrate that maternal *Axl* knockout was the cause of abnormal maternal-fetal crosstalk of PE syndrome.

**Figure 3.**
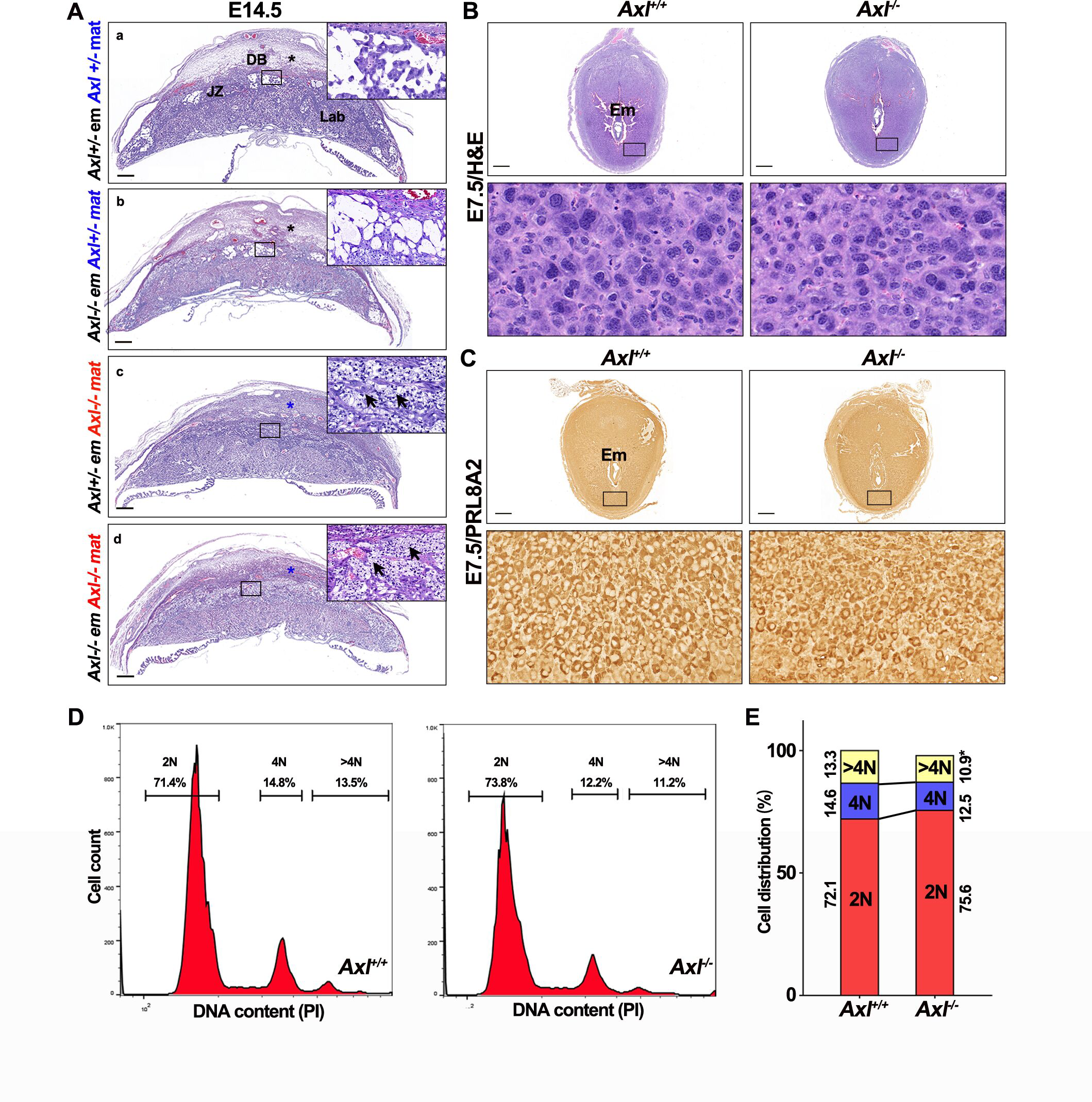
Maternal *Axl* knockout leads to abnormal interface in *Axl^−/−^* mice. (A) Utero-placental interface structure of different mating strategies by H&E staining. Black arrows point to foci of glycogen deposition. Black asterisks mark the loosened decidual zone while blue asterisks point the compacted decidua. (B) The histology of E7.5 implantation sites and polyploid stromal cells in *Axl^+/+^* and *Axl^−/−^* females. (C) The expression of decidual marker PRL8A2 at *Axl^+/+^* and *Axl^−/−^*implantation sites on E7.5. (D) DNA content quantification of decidual cells by FACS from *Axl^+/+^* and *Axl^−/−^* mice on E7.5. (E) Cellular distribution for 2 N, 4 N, and >4 N populations of stromal cells at *Axl^+/+^*(n = 3 animals) and *Axl^−/−^* (n = 3 animals) implantation sites on E7.5. Data represent the mean ± SEM. Scale bars: 0.5 mm. *, *p* < 0.05. DB, decidual basalis; JZ, junctional zone; Lab, labyrinth; Em, embryo.

Previously studies have demonstrated that, rather than passively undergoing modification directed by invasive trophoblasts, mouse uterine stromal cells initiate decidualization and transform into secretory decidual cells when embryos implant before they can function in modulating the maternal-fetal interface (Menkhorst, Van Sinderen et al., 2019). To screen for abnormalities in *Axl^−/−^*decidual cells, we examined stromal decidualization using early pregnant uteri. On E7.5, H&E staining showed that the implantation sites in *Axl^−/−^* mice were morphologically normal, and the decidual cells were transformed into multinucleated secretory cells apparently normally (Figure 3B). We further examined the expression of decidualization markers. PRL8A2 is a well-known marker of mouse decidualization, and immunohistochemistry (IHC) results showed no obvious difference in the expression of PRL8A2 between *Axl^+/+^* and *Axl^−/−^* uterine decidua (Figure 3C). However, the size of decidual cells was manifestly smaller in the *Axl^−/−^* mice than that in their *Axl^+/+^*counterparts (Figure 3B and 3C). Flow cytometric analysis of DNA content revealed a concomitant decrease in >4N cells in *Axl^−/−^* decidua on E7.5 (Figure 3D and 3E), indicating insufficient decidualization in the *Axl^−/−^*mice and suggesting molecular abnormalities.

### Down-regulated *Corin* via STAT3 pathway contributes to the defective decidualization in *Axl^−/−^* mice

To explore the molecular mechanisms underlying the defects in *Axl^−/−^*decidua, we employed artificially induced decidualization by oil injection, and observed induction of multinucleated decidual cells in both *Axl^+/+^*and *Axl^−/−^* mice (Figure 4A and 4B). We further explored the molecules in *Axl^−/−^*decidua by performing RNA-seq of the induced deciduoma samples considering the above *in vivo* data of *Axl^−/−^* mice. A heatmap representing the clustering of differentially expressed genes was established (Figure 4C). Comparing the transcriptomes of the decidualized cells in the two groups revealed 850 dysregulated genes (˃ twofold), including 423 downregulated genes and 327 upregulated genes in *Axl^−/−^* deciduoma compared to *Axl^+/+^* counterparts shown in Figure 4D. We noticed that *Corin*, the gene encoding an ANP converting enzyme, was downregulated in *Axl^−/−^* deciduoma (Figure 4D). A previous study showed that both CORIN and ANP play essential roles at maternal-fetal interface, and their defects contribute to PE by impairing trophoblast invasion and SpA remodeling (Cui, Wang et al., 2012). We therefore confirmed differential expression of angiogenesis (*Angptl1*, *Ada*) and renin-angiotensin system (RAAS)-related genes (*Corin, Ren2*) by qPCR analysis (Figure 4E). These data indicate that the downregulated AXL-CORIN signaling contributes to the defective decidualization in *Axl^−/−^*decidua.

**Figure 4.**
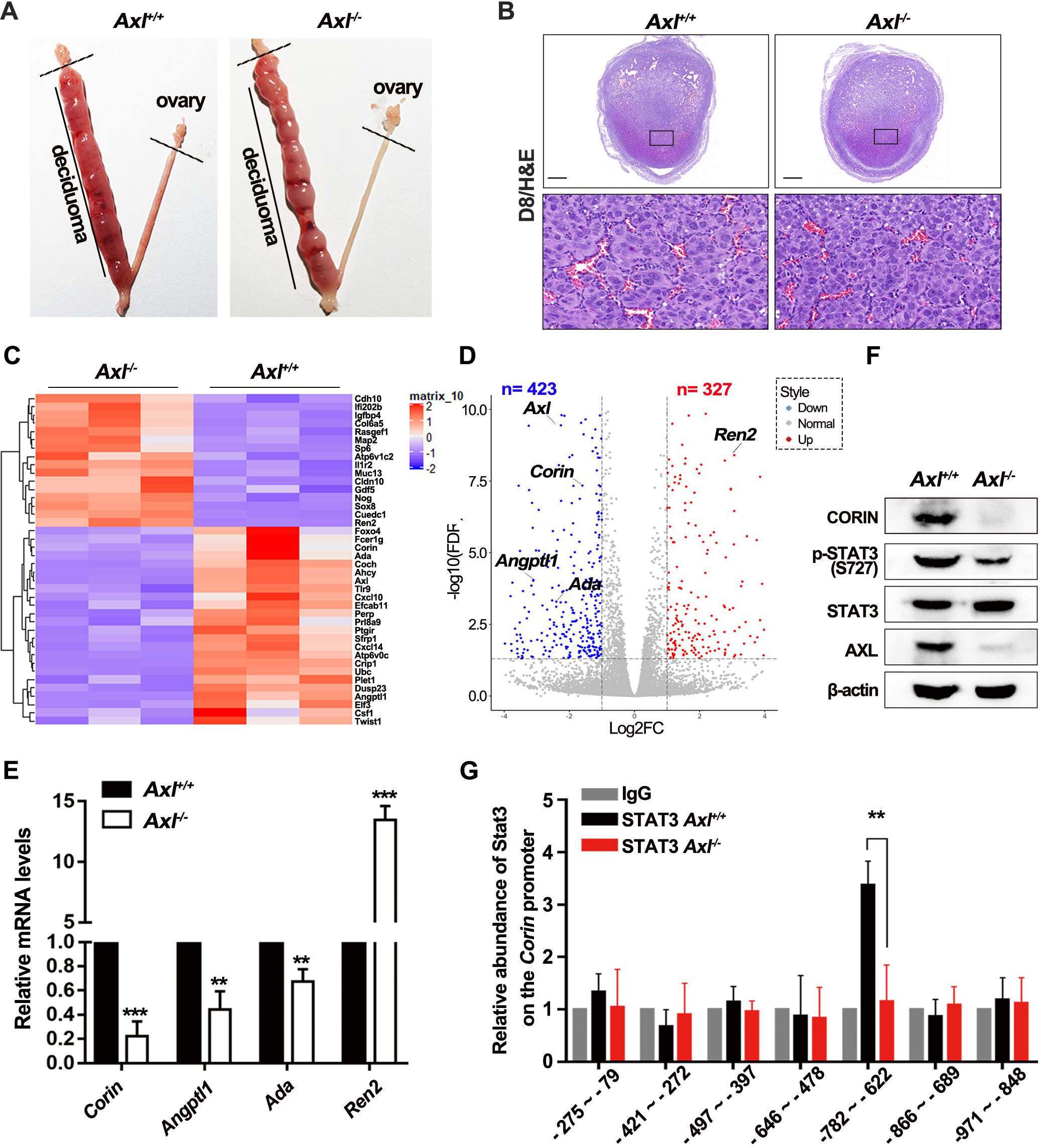
Decreased *Corin* expression in *Axl^−/−^* decidua. (A) Gross morphology of deciduoma at E8.5 of pseudopregnancy after artificial decidualization is induced in *Axl^+/+^*and *Axl^−/−^* mice. (B) H&E staining of deciduoma sections from the oil-instilled left horn of the uterus showed a similar area in *Axl^+/+^* and *Axl^−/−^* pseudo-pregnant mice after 96 hours after the oil injection. (C) Heatmap of genes differentially expression of E8.5 oil-induced *Axl^−/−^* deciduoma compared with *Axl^+/+^* ones. Red to black to blue indicates a gradient of high to low expression. (D) Volcano plot of the expression of altered genes. Genes that are differentially expressed at FDR < 0.05 and log FC > 1 are highlighted and labeled. (E) Real-time qPCR validation of 5 genes selected from the Volcano plot. The error bars represent standard error of mean (SEM). (F) Western blot analysis of CORIN and STAT3 activation level in both *Axl^+/+^* and *Axl^−/−^* deciduoma. β-actin was used as the control. (G) The binding of STAT3 to *Corin* promoter was decreased in *Axl^−/−^*mice. ChIP assays were performed using anti-STAT3 and control IgG antibodies and fold enrichment of the indicated regions of CORIN promoter was examined using real-time PCR in isolated *Axl^+/+^* and *Axl^−/−^* primary decidual cells. All ChIP samples were normalized to intergenic input control. Scale bars: 0.5 mm. **, *p* < 0.01; ***, *p* < 0.001.

We subsequently explored the molecular mechanism by which *Axl* knockout leads to *Corin* downregulation. Immunoblotting of decidual tissue of *Axl^−/−^*mice showed dramatically decreased expression of CORIN relative to the control decidua. Activation of the JAK-STAT3 pathway, to be canonical downstream target of AXL was impaired, compared to *Axl^+/+^* decidua (Figure 4F). These data demonstrate that *Axl* knockout led to the downregulation of CORIN via the JAK-STAT3 pathway in the decidua. Then we studied the relationship between *Corin* and JAK-STAT3 by ChIP-qPCR with a STAT3 antibody on decidual tissue samples. We observed that, relative to the STAT3 binding peak for a specific region (−782bp ∼ −622bp) of the *Corin* in *Axl^+/+^*decidua, the extent of binding between these two factors was significantly reduced in *Axl^−/−^* decidua (Figure 4G). These data demonstrate that *Axl* knockout downregulates CORIN via the JAK-STAT3 pathway in the decidua and that this impaired signaling contributes to defective decidualization, causing the poor communication between maternal decidua and fetal trophoblasts.

### ANP rescued the PE symptoms in *Axl* knockout mice by promoting trophoblast invasion

Since the cardiac protease, CORIN, activates ANP, a cardiac hormone that regulates blood pressure (Chen, Cao et al., 2015, Cui et al., 2012), the downregulation of CORIN in *Axl^−/−^*decidua would be expected to deplete ANP abundance. To test this hypothesis, we injected *Axl^−/−^* mice daily with ANP (2 μg/0.1 ml/ per mouse) or vehicle control (saline) by intraperitoneal injection from E8.5 to E18.5 to see if it can rescue the PE symptoms. We noticed that, compared to the continuously increased SBP before E16.5 observed in the vehicle control *Axl^−/−^* mice, both the vehicle-treated *Axl^+/+^* mice and the ANP-treated *Axl^−/−^*mice had normotensive blood pressure (Figure 5A). The ANP treatment also rescued the proteinuria symptom of the *Axl^−/−^* mice (Figure 5B). These data demonstrated that ANP can indeed rescue the PE symptoms in *Axl^−/−^* mice.

**Figure 5.**
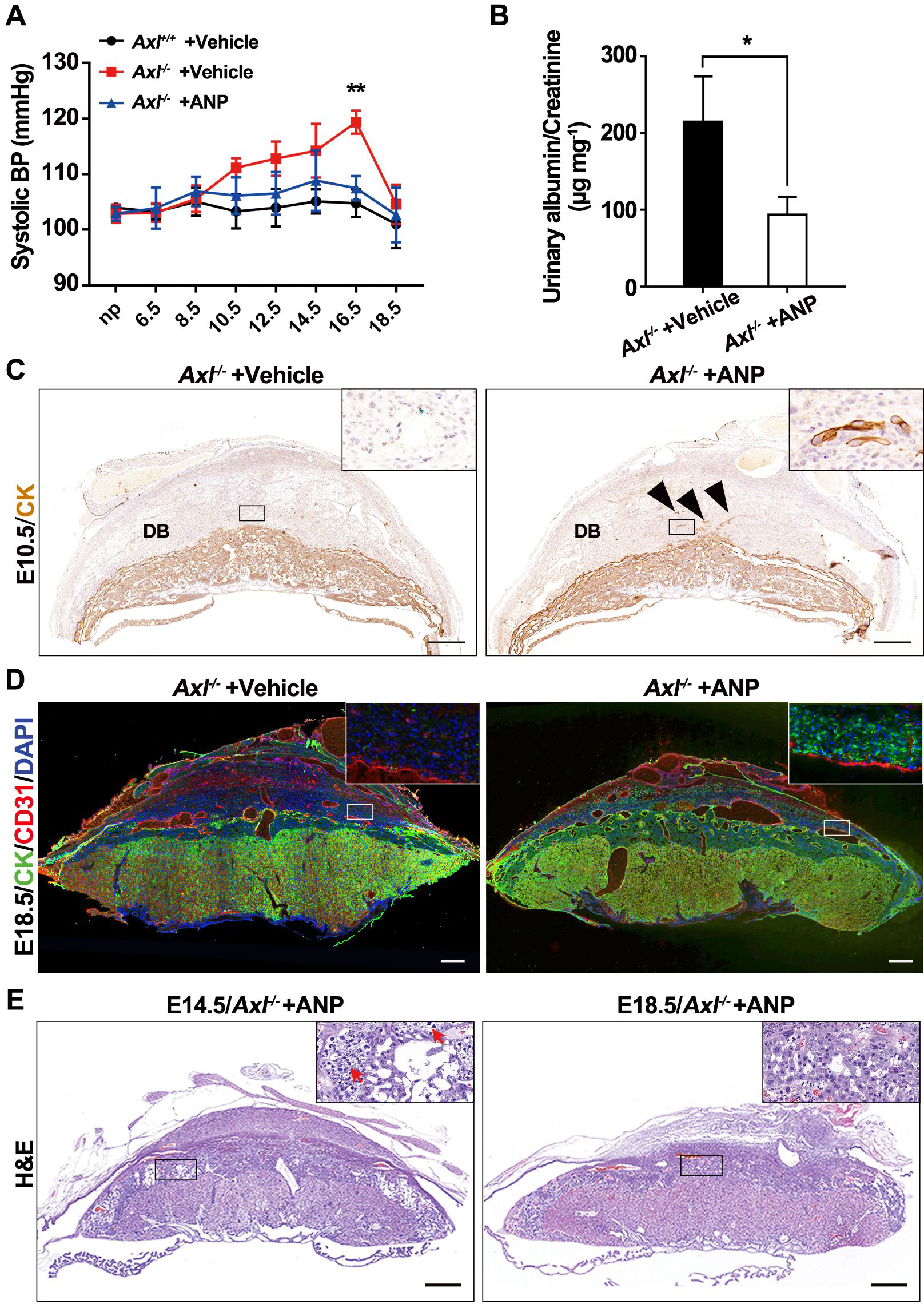
ANP treatment rescues PE symptoms in *Axl^−/−^* mice. (A) Mean systolic blood pressure increased in *Axl^−/−^* mice treated with saline but not in those with ANP (2 μg/0.1 ml/day per mice). (B) Late gestational urine Albumin/Creatine in *Axl^−/−^* mice treated with saline or ANP. n=3 per group. (C) Immunohistochemical analysis of trophoblast cell marker (cytokeratin, CK) at *Axl^−/−^* interfaces with vehicle (saline) or ANP treatment at E10.5. Black triangles marked invasive trophoblast cells. (D) Photomicrographs of trophoblast cell marker (CK) and blood vessel endothelial marker (CD31) at E18.5 maternal-fetal interface of *Axl^−/−^*mice treated with vehicle or ANP. (E) H&E staining of *Axl^−/−^*mice treated with ANP at E14.5 and E18.5. (F) Quantification of the average area of the junctional zone and labyrinth of placentas from different groups. Scale bars, 0.5 mm. Data are means ± SEM. *, *p* < 0.05; **, *p* < 0.01. DB, decidual basalis.

To establish whether ANP can relieve the *Axl^−/−^* mice placental phenotype in addition to alleviating the blood pressure and proteinuria symptoms, we explored the uteroplacental interface of *Axl^−/−^* mice by IHC, immunofluorescence and H&E staining. The IHC results demonstrated that ANP promoted trophoblast invasion into the decidua at E10.5 in *Axl^−/−^*mice, consistent with the invasion seen in the *Axl^+/+^*line (Figure 5C). In addition, CD31 and cytokeratin immunofluorescence staining showed that SpA lumen was enlarged and the numbers of invaded trophoblasts increased in *Axl^−/−^* decidua treated with ANP at E18.5 (Figure 5D). These data demonstrate that ANP enhanced trophoblast invasion and SpA remodeling, thus achieving better maternal and fetal interaction. Moreover, when we examined the placenta by H&E staining, we can still find glycogen deposition (red arrows) in the junctional zone of *Axl^−/−^*mice on E14.5 after ANP treatment; and it was greatly relieved at E18.5 (Figure 5E), indicating the status of *Axl^−/−^* mice is continuously improved.

### AXL expression is decreased in the decidual tissues of preeclamptic women

Decidual tissue sections of patients with severe PE and normal pregnancy (Table S1) were assessed with IHC against AXL (Figure 6A). Using VIMENTIN as a marker for decidual cells quantified with Image-Pro plus 6.0, we observed decreased AXL expression in the severe PE group compared to the normal pregnancy group (Figure 6A and 6B). In addition, we examined the mRNA for *AXL* by qPCR, and found its abundance to be significantly downregulated in severe PE group compared to the normal group (Figure 6C). Western blotting of the decidual tissue indicated that the AXL protein level was significantly decreased in the severe PE group compared to the normal group (Figure 6D and 6E). These data indicate that decidual AXL expression is reduced in these PE patients.

**Figure 6.**
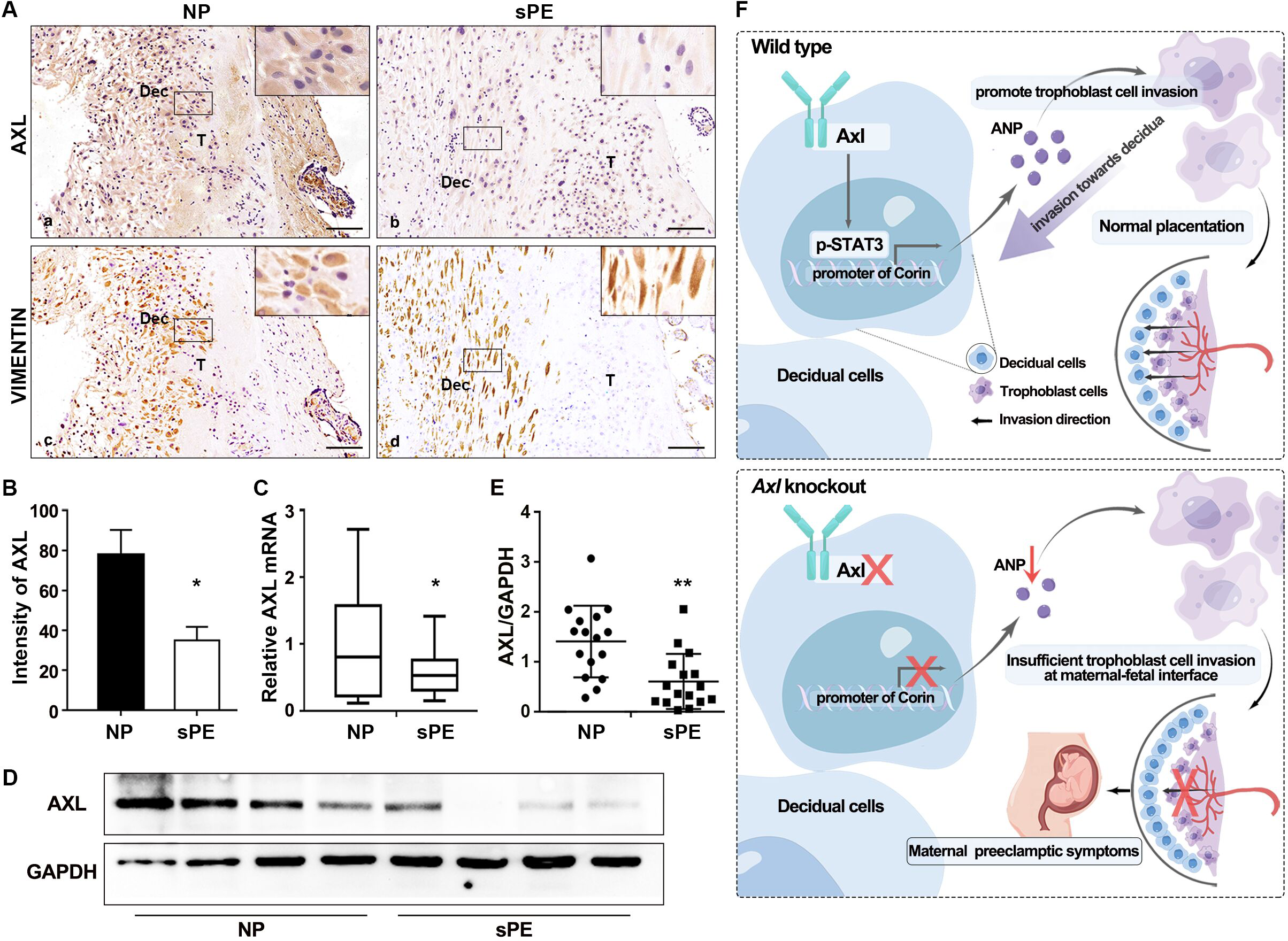
The expression of AXL in decidua of normal pregnancy and sPE patients. (A and B) Representative images and quantification of AXL intensity in the decidual tissue of normal pregnancy (NP) and severe preeclampsia (sPE). (a, b) AXL; (c, d) Vimentin. Dec, decidual cell-enriched area; T, trophoblast cell-enriched area. (C) Relative expression of AXL mRNA to GAPDH in utero-fetal interfaces of NP and sPE. (D) Western blotting showed the protein level of AXL was decreased in sPE utero-fetal interfaces. GAPDH was used as the loading control. (E) Western blotting results were quantified and shown in bar graphs (NP vs sPE, 17 vs 16). (F) Pathway model in which *Axl* knockout leads to insufficient placental trophoblast invasion through STAT3-CORIN-ANP signaling, thus causing maternal preeclamptic symptoms. Our study highlights the role of aberrant interactions between the maternal decidua and fetal placenta in the pathogenesis of preeclampsia. Data are presented as means ± SD. Scale bars, 0.2 mm. *, *p* < 0.05; **, *p* < 0.01.

## Discussion

In this study, we show that *Axl^−/−^* pregnant mice exhibit the typical PE symptoms, including elevated SBP, proteinuria with abnormal glomerular structures, reduced placental and fetal weight. ANP treatment increased trophoblast invasion, SpA remodeling and rescued glycogen deposition at the maternal-fetal interface of *Axl*^−/−^ females. Further investigation verified that maternal decidua contributed to the PE symptoms, and although the decidualization was morphologically normal, the molecular expression and regulation were defective. In addition, *Axl*^−/−^ the decidual basalis accumulated more extracellular matrix and was more compact. Dysregulated JAK-STAT3 in *Axl*^−/−^ decidual tissue reduced CORIN expression. Furthermore, AXL expression was significantly decreased in PE patients, indicating AXL-CORIN-ANP plays critical role in the pathogenesis of PE.

Glomerular endotheliosis is characterized by enlarged glomeruli that lack blood cells due to the swollen endothelial cells and occluded capillary lumina (Stillman & Karumanchi, 2007). In our study, analysis of kidney sections revealed that in *Axl^−/−^*pregnant females display abnormal glomerular structures, including narrow capillary lumen, thickened basement membrane, and shorted foot process, which decreases glomerular filtration and results in proteinuria due to endothelial destruction. Our findings are similar to a previous report that renal tissue in PE exhibits hallmarks of glomerular endotheliosis (Phipps, Thadhani et al., 2019).

By comparing *Axl^+/+^* and *Axl^−/−^* interfaces, we found that the decidua of *Axl^−/−^* mice is more compact and has non-degraded extracellular matrix. Among the many roles played by the extracellular matrix are regulating the formation of new vessel sprouts and provision of the necessary contacts between endothelial cells and surrounding tissue, thus preventing vessels from collapsing (Carmeliet, 2003). When vascular cells migrate to form new sprouts, this matrix network is proteolytically degraded and its composition is altered. Reduction of the extracellular matrix during vessel sprouting is therefore essential for SpA remodeling and angiogenesis. We conclude that the persistence of non-degraded extracellular matrix in the decidua of *Axl^−/−^* mice contributes to the PE symptoms observed in this model.

The maternal-fetal interfaces that resulted from cross mating and embryo transfer strategies employing different genotypes demonstrated that *Axl^−/−^* maternal decidua is the cause of the abnormality. Our trials comparing gestational decidua and artificial oil-induced decidua show decidualization occurs morphologically normally in the *Axl^−/−^*mouse relative to the control counterpart. The principal difference is that the decidua is more compact. Our subsequent RNA-seq, real-time qPCR, and Western blot analysis of oil-induced deciduoma revealed the downregulated expression of CORIN in decidual tissue in the *Axl^−/−^*mouse, consistent with women complicated with PE (Cui et al., 2012). The downstream consequence of AXL signaling is decreased activation of STAT3 pathway. There is a pronounced reduction in activation of this pathway in *Axl^−/−^* decidua. Our further studies using ChIP-qPCR found that STAT3 can directly bind to the *Corin* promoter, and the downregulation of the STAT3 pathway in *Axl^−/−^* decidua inactivated *Corin* promoter. The deficiency of *Corin* or its product, ANP, impairs SpA remodeling, causing gestational hypertension and proteinuria, the major PE symptoms (Armstrong, Tse et al., 2014, Cui et al., 2012). Moreover, ANP has been reported in promoting trophoblast invasion in a TRAIL-dependent way and decidualization (Zhang, Li et al., 2021), which helps us to prove that ANP rescue the symptoms of *Axl^−/−^* pregnant mice. Our finding of disturbance of the AXL-STAT3-CORIN pathway sheds light on how ANP could rescue the *Axl^−/−^* symptoms. The reduced expression of *Corin* in *Axl^−/−^* females due to JAK-STAT3 dysregulation in the uterine decidua contributes to PE though their decidualization is morphologically not affected.

ANP has been shown to facilitate trophoblast cell invasion in vitro (Cui et al., 2012, Wang, Wang et al., 2020). In HTR8/SVneo cells, a cell line of human invasive cytotrophoblasts, knockdown of the ANP receptor, *NPR-A* with siRNA impaired its invasive ability, but had no effect on its proliferation or frequency of apoptosis (Tan, Lin et al., 2019). These findings are concordant with our observations in the mouse decidua. In addition, there are several other decidua-derived factors involved. Previous reports have also shown that the invasion of trophoblast cells into the uterus is regulated in situ by a number of locally produced molecules, such as transforming growth factor-beta and decorin, essential for maintaining healthy maternal-fetal homeostasis (12,34). Collectively, our study provides convincing evidence that the decidua-derived cascade of AXL-CORIN-ANP signaling is essential to proper trophoblast invasion to ensuring well-modified vascular connections between the mother and the fetus. In contrast, even decidual defects induced by AXL-deficiency impair decidualization, contribute to this life-threatening disease, underlining the essential role of decidual signaling in regulating pregnancy.

## Materials and Methods

### Animal care and genotyping

*Axl*^−/−^ mice on C57BL/6J background were purchased from Jackson Laboratory (ME, USA), and genotyping was performed as previously reported (Lu et al., 1999). *Axl*^−/−^ mice were crossed to obtain the *Axl*^+/-^ mice, and the *Axl*^+/+^ littermates were used as controls. All mice were housed under controlled temperature (22 °C±3 °C) and light conditions (14 h light, 10 h darkness; lights on at 07:00 AM), and allowed free access to chow and water at the Experimental Animal Center of Shandong Normal University. All animal experiments were approved by the Animal Ethics Committee of Shandong Normal University.

### Analysis of pregnancy and the artificial decidualization model

Two-month-old *Axl*^+/+^ and *Axl*^−/−^ female mice were mated with fertile or vasectomized males of the same strain to induce pregnancy or pseudo-pregnancy, respectively. The morning of finding a vaginal plug was designated embryonic day 0.5 of pregnancy (E0.5). To induce artificial decidualization, one uterine horn of pseudo-pregnant mice was infused with sesame oil (25 μl) on E3.5; the non-infused contralateral horn was taken as a control. The tissue of infused and non-infused uterine horns was collected 4 days after the oil infusion. A section of each uterus was snap frozen in liquid nitrogen for subsequent RNA-seq analysis. A minimum of three mice were used for every individual experiment in each mouse model.

### Blastocyst transfer

*Axl^+/+^* and *Axl^−/−^* female mice (donor mice) were injected with 7.5 U pregnant mare serum gonadotropin (Tianjin Laboratory Animal Center, Tianjin, China) and 48 h later with 7.5 U human chorionic gonadotropin (Tianjin Laboratory Animal Center, Tianjin, China) to induce ovulation; injected female mice then were mated. Meanwhile, *Axl^+/+^*and *Axl^−/−^* female mice (receiver mice) were mated with *Axl^+/+^*vasectomized males to induce pseudopregnancy. At E3.5, donor females were killed by cervical dislocation, and blastocysts were collected and transferred into the uterine horn of receiver mice.

### Flow cytometry

E7.5 decidual cells were digested, centrifuged, and suspended in PBS; 1 ml of cold 80% ethanol was added dropwise with continued gentle vertexing and incubated for 30 min on ice before staining. Cell pellets were suspended in staining solution (5 mg/ml Propidium Iodide and 2 mg/ml DNase-free RNase A in PBS) for 30 min at 37°C in the dark before flow cytometry was performed using Beckman Cytoflex. Flow cytometry data were analyzed using FlowJo software (v10, Tree star). Fluorescence-activated cell sorting (FACS) intensity values for SSC-A and FSC-A of were used for loose gates. The intensity values of FSC-A and FSC-H by FACS were used to gate individual cells. Polyploid cells were differentiated by FSC-A intensity values proportional to cell size and propidium iodide staining intensity values used to measure DNA content. The experiments were repeated three times.

### Blood pressure monitoring

The blood pressure was measured by a noninvasive blood pressure system for mice and rats (Softron Biotechnology, Beijing, China), as described previously (Wei, Li et al., 2007). The investigators were blinded during the measuring of the blood pressure. SBP was calculated from three consecutive recordings. Data presented were from continuous recording of at least 6 h per day (10:00 to 16:00).

### Urinary albumin/creatinine measurements

Urine samples were collected for 72 h in metabolic cages from E14.5 to E17.5 in the presence of a mix of proteinase inhibitors (Roche, Hvidovre, Denmark). Urine creatinine concentrations were determined using the Creatinine Companion kit (Cayman chemical company, Ann Arbor, MI), and albumin concentrations were determined using a mouse albumin ELISA quantification kit (Bethyl Labs, Montgomery, TX).

### Uterine artery Doppler ultrasound

More than three pregnant mice of both *Axl^+/+^* and *Axl^−/−^* genotype on E14.5 were imaged transcutaneously using Doppler ultrasound and a 30- or 40-MHz transducer operating at 30 frames/s (model Vevo 660, VisualSonics, Toronto, Canada). Studies were performed between 1 PM and 5 PM. The high-pass filter was set at 6 Hz, and the pulsed repetition frequency was set between 4 and 48 kHz, to detect low to high blood flow velocities, respectively. A 0.2- to 0.5-mm pulsed Doppler gate was used, and the angle between the Doppler beam and the vessel was adjusted from 0 to 60 degrees to provide the best alignment. Waveforms were saved for later offline analysis. The duration of anesthesia was limited to ∼1 h. The resistance index (RI) and the pulsatility index of the uterine arteries were calculated.

### Transmission electron microscopy (TEM)

For TEM analysis, E14.5 *Axl^+/+^* and *Axl^−/−^*kidneys were fixed with 2.5% glutaraldehyde at room temperature for 2-3 h, placed at 4°C. Then were fixed in 1% osmic acid for 1.5 h in the dark. The samples were then graded dehydrated in ethanol and infiltrated with acetone: resin=1:1 for 1 h, acetone: resin=1:2 overnight, and finally infiltrated with resin twice for 4 h each time. After embedding in resin, the samples were trimmed manually and sections were cut (65 nm) using a Leica UC7 Ultramicrotome (Hitachi Ltd, Hitachi, Japan). The sections were stained with 2% uranyl acetate solution for 25 min, washed with water, and then stained with lead citrate for 7 min. The sections were subsequently baked under infrared light for 10 min and then observed under an 80 KV transmission electron microscope (Hitachi HT-7800, Hitachi Ltd, Hitachi, Japan), and photographs were taken at different magnifications.

### Immunofluorescence

Pregnant uteri were snap frozen in liquid nitrogen and then cryosectioned at 10 μm in at −20°C. The slides were fixed with ice acetone for half an hour, cleaned with PBS, and then permeabilized for 10 min in 0.1% Triton X-100. After blocking with 10% normal goat serum in PBS for 1 h at room temperature, the sections were incubated with primary antibodies in blocking solution overnight at 4 °C. Specific secondary antibodies were used to detect the antigen and 4’,6’-diamidino-2-phenylindole (DAPI) was applied to identify the cell nucleus. All samples were imaged using a Leica confocal microscope (TCS SP8 MP, Leica Microsystems, Wetzlar, Germany).

### H&E and immunohistochemistry (IHC)

Isolated tissues were fixed overnight in 4% paraformaldehyde (PFA) in PBS, dehydrated using an ethanol series, embedded in paraffin and sectioned (5 μm). For H&E staining, the tissues were deparaffinized in xylene, rehydrated through a series of ethanols, and stained with H&E. For IHC, deparaffinized sections were subjected to antigen retrieval by boiling for 1 h in Tris/EDTA (pH 9.0) and cooling to room temperature, followed by peroxidase blocking for 10 min at room temperature. The blocking and antibodies were used for the immunofluorescence process described above. Sections were then incubated overnight at 4 °C with the primary antibodies, then incubated for 20 min at room temperature with horseradish peroxidase-labeled goat anti-rabbit IgG (Zhongshan Company, Beijing, China), and rinsed with PBS. The antibody complexes were detected using DAB reagent according to the manufacturer’s instructions (Zhongshan Company, Beijing, China). Haematoxylin staining was then performed as per standard protocols. Antibodies used are listed in Table S2. All samples were imaged on a Pannoramic Scan (3Dhistech Company, Budapest, Hungary). The staining quantification was performed using Image-Pro plus 6.0 image analysis software (Media Cybernetics, Silver Spring, MD, USA), for each group, total of 30 fields from 10 different women (each woman 3 fields of view) were photographed and examined.

### Periodic acid schiff (PAS) staining

To detect the corpora amylacea, paraffin sections were incubated in 0.5% periodic acid (Solarbio Company, Beijing, China) for 5 min at room temperature, followed by washing in tap water for 1 min. Then the sections were incubated in Schiff reagent (Solarbio Company, Beijing, China) for 10 min, followed by washing in tap water for 10 min. After the sections were rinsed in PBS for 10 min, the nuclei were stained with hematoxylin.

### SDS-PAGE and western blotting

The lysates containing 30 μg of protein were loaded on a 10% SDS-polyacrylamide gel, separated with 1xTris-Glycine running buffer, and then transferred to PVDF membranes (Millipore, MA, USA) in a wet electroblotting system. PVDF membranes were blocked for 1 h in 5% nonfat milk in TBST, then incubated overnight at 4℃with antibodies. After washing in TBST, the membranes were incubated with secondary antibody (ZSGB-BIO, Beijing, China) in TBST, and then detected with the SuperSignal chemiluminescent detection system (Thermo Scientific, CA, USA). Antibodies used are listed in Table S2. GAPDH served as an internal loading control.

### RNA isolation and quantitative polymerase chain reaction (PCR)

Total RNA was isolated with the TRIzol Reagent (Invitrogen, CA, USA) and quantified with a NanoDrop 2000 (Thermo Scientific, CA, USA). An aliquot of 1 μg RNA was used to synthesize cDNA. Expression levels of the genes were validated by quantitative real-time PCR analysis with SYBR Green (Takara Bio, CA, USA). Cycling parameters were 95°C for 20 seconds, followed by 40 cycles of 95°C for 3 seconds and 60°C for 30 seconds. The efficiency of all SYBR green primer sets was determined and validated using serial dilutions to generate a 5-point curve. The primers used for test genes are listed in Supplementary Table S3.

### Construction of RNA sequencing libraries and sequencing

Total RNA was extracted from the samples using Trizol reagent (Invitrogen). The RNA quality was detected with Agilent 2200 and stored at −80°C. The RNA integrity number (RIN) > 7.0 was used for the construction of complementary DNA (cDNA) libraries. A cDNA Library was constructed for each RNA sample using the TruSeq Strand mRNA Library Prep Kit (Illumina, Inc.) according to the manufacturer’s instructions. In brief: mRNA was purified from 1ug of total RNA using oligomeric (dT) magnetic beads and cleaved into 200-600 bp RNA fragments using divalent cations at 85°C for 6 min. The cleaved RNA fragments were used for the synthesis of the first and second strands. The cDNA fragment was end-repaired and attached to the index adapter. The ligated cDNA product was purified and treated with uracil DNA glycosylase to remove the second strand cDNA. The purified first-strand cDNA was enriched by PCR to establish a cDNA library. The libraries were quality-controlled with Agilent 2200 and sequenced with NovaSeq 6000 on a 150 bp paired-end run.

### RNA-seq data analysis

Clean reads are obtained from raw reads by removing adapter sequences and low-quality reads prior to read mapping. The clean reads were then aligned to the mouse genome (mm10, NCBI) using Hisat2 (Kim, Langmead et al., 2015). HTseq (Anders, Pyl et al., 2015) was used to obtain gene counts and RPKM method was used to determine gene expression. DESeq2 algorithm (Love, Huber et al., 2014) was used to filter the differentially expressed genes. After significant analysis, P-value and FDR analysis (Benjamini, Drai et al., 2001) were subjected to the following criteria: i) Fold Change>2 or < 0.5; ii), P-value< 0.05, FDR<0.05. Gene ontology (GO) analysis was performed to elucidate the biological implications of the differentially expressed genes (Ashburner, Ball et al., 2000). Fisher’s exact test was applied to identify the significant GO categories (P-value < 0.05).

### ChIP‒qPCR

STAT3 binding to *Corin* promotor was assessed by ChIP assay using *Axl^+/+^* and *Axl^−/−^* decidua. ChIP analysis was performed as previously described (Zhang, Meng et al., 2019). In brief, the isolated tissues were fixed with formaldehyde for 10 min at room temperature. The chromatin was sonicated using the Bioruptor® Pico sonication device (Diagenode Diagnostics, Belgium) to fragment the DNA-protein complexes. This supernatant was then incubated overnight at 4°C with STAT3 antibody, followed by protein A/G beads to pulldown the complex. After several washes, the resulting protein/DNA complexes were subjected to cross-link reversal and DNA extraction. Specific primers were used to detect immunoprecipitated chromatin fragments, as well as input chromatin, using qPCR assay (the primers used are listed in Table S3).

### *In vivo* treatment with ANP

Pregnant *Axl^−/−^* mice were treated with ANP (2 μg/0.1 ml/ per mouse) or vehicle control (saline) intraperitoneal injection daily from E8.5 to E18.5. The blood pressure was measured every other day since E6.5. Mice were sacrificed at E18.5 to collected utero-placental tissues.

### Human subjects

Sixteen women with severe PE (sPE) and 17 women with normal pregnancies (NP) were recruited in the Department of Obstetrics and Gynecology, Ren Ji Hospital, School of Medicine, Shanghai Jiao Tong University between May 2019 and July 2021. sPE is defined as systolic blood pressure ≥ 160 mmHg on at least 2 occasions 6 h apart or with proteinuria >3+ protein on dip stick after 20 weeks of gestation. Only caesarean pregnancies were included and none of the included mothers were in labor prior to caesarean section. Multiparous pregnancies, renal disease, maternal diabetes, chromosomal aberrations, fetal and placental structural abnormalities were excluded from our study. Decidual tissues were obtained by scrubbing the uterine wall at the site of the placental bed with a gauze after the placenta was delivered. The decidua was washed with sterilized water to remove blood, snap frozen in liquid nitrogen and stored at −80 °C until use. The study was approved by the Renji Hospital Research and Ethics Committees. Informed consent was obtained from all participants before the collection of decidual samples.

### Statistical analysis

GraphPad Prism 8.0 software (GraphPad Software, San Diego) was used for statistical analysis. Student’s unpaired two tailed t-tests were used for statistical analyses. *P* < 0.05 was considered statistically significant. Sample sizes were selected based on current and previous experiments and no statistical method was applied to predetermine sample size. All experiments were repeated at least three times.

## Acknowledgements

This study was supported by the National Key R&D Program of China (2019YFA0802600) and the National Natural Science Foundation of China (32170863 and 31871512) to C. Zhang. Support was also received from grants from the Shanghai Commission of Science and Technology (20DZ2270900) and Open Project of Shandong Provincial Key Laboratory of Reproductive Medicine (SDKL2017018).

## Author contributions

C. Zhou., Y.Z., Z.-J. C. and C. Zhang planned mouse experiments, which were carried out by C. Zhou., Y.Z., L. Z., X. L., J. Y. and C. Zhang. C. Zhou and C. Zhang wrote the manuscript. C. Zhou and C. Zhang planned and executed human decidua experiments and edited the manuscript. Z.-J. C. edited the manuscript

---

**Figure S1.**
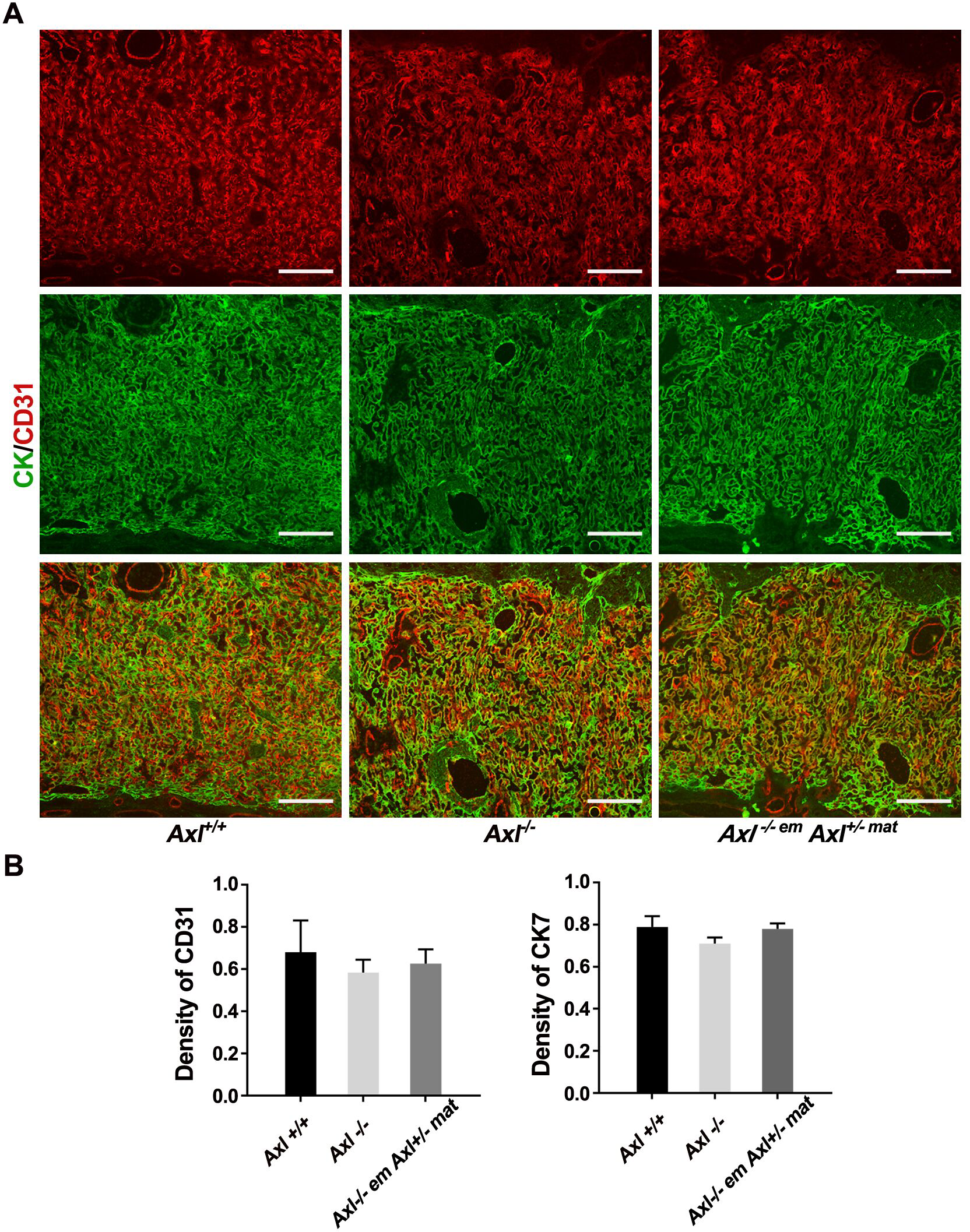
AXL deletion impaired junction area but not labyrinth. (A) Photomicrographs of trophoblast cell marker (cytokeratin, CK) and blood vessel endothelial marker (CD31) at E14.5 maternal-fetal interfaces of different groups. (B) Quantification of vascular and trophoblastic density using CD31 and CK, respectively. Scale bars, 0.2 mm.

**Figure S2.**
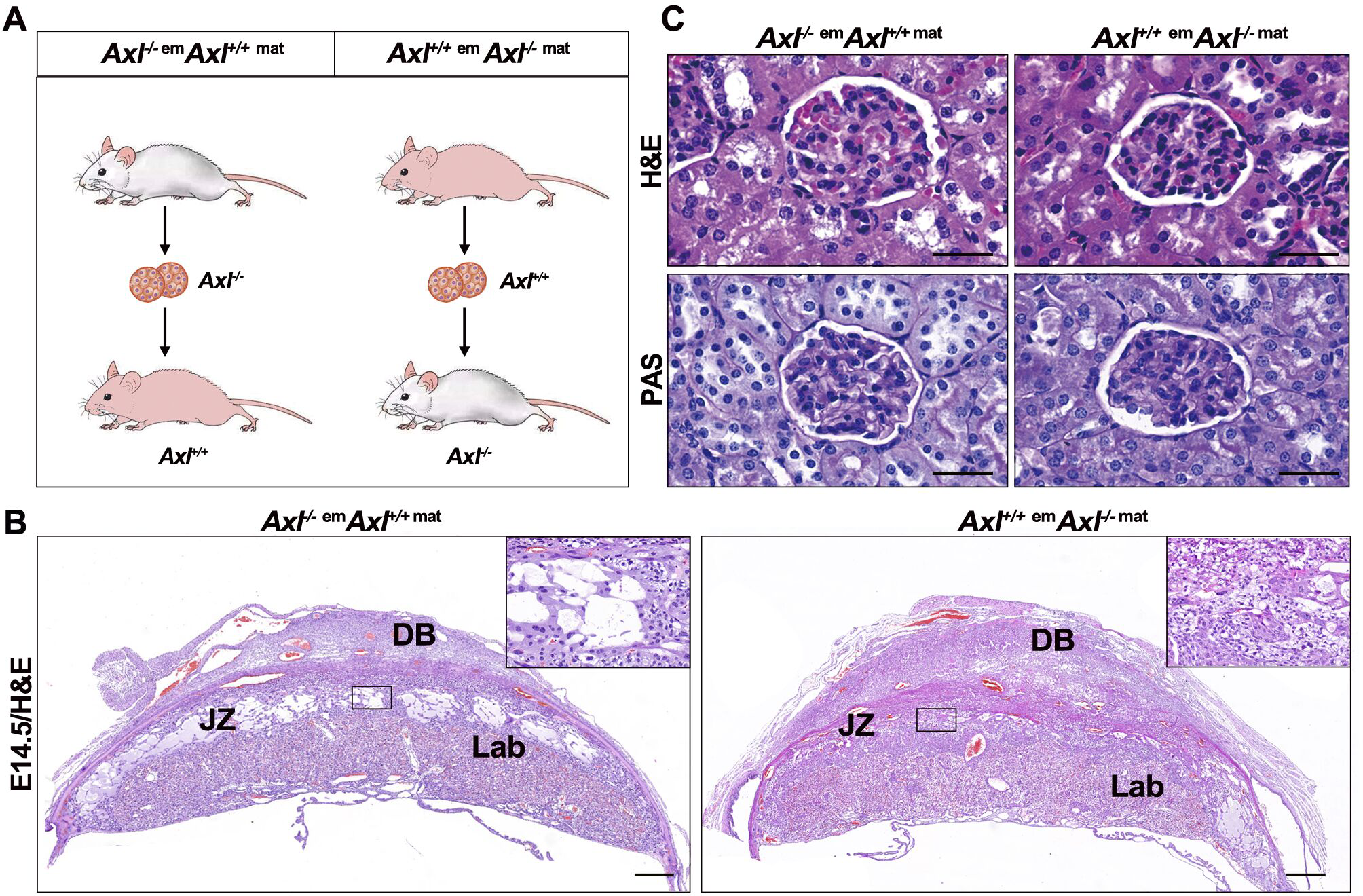
Embryo transferring between *Axl^+/+^* and *Axl^−/−^* mice. (A) Sketch map of embryo transfer. (B) The placenta from *Axl^−/−^*mice carrying *Axl^+/+^* fetus showed large confluent areas of glycogen deposition, replacing much of the JZ at E18.5. DB, decidual basalis; JZ, junctional zone; Lab, labyrinth. Scale bars, 0.5 mm. (C) Comparison of H&E and PAS staining of the renal tissues in embryo transferred mice. Results indicate increased extracellular matrixes in *Axl^−/−^* females with *Axl^+/+^* embryo. Scale bars, 50 μm.

**Table S1.** The clinical characteristics of study subjects in the sPE and Normal Pregnancy groups.

| Variables | sPE (n=16) | NC (n=17) | P value |
| --- | --- | --- | --- |
| Age(years) | 32.19±2.903 | 29.65±3.334 | 0.5969 |
| Gestational age at delivery (weeks) | 37.19±2.563 | 38.82±3.588 | <0.01 |
| SBP (mmHg) | 150.5±11.237 | 119±9.062 | <0.01 |
| DBP (mmHg) | 94.06±6.527 | 75.94±8.863 | <0.01 |
| Birth weight (g) | 3043.75±457.038 | 3547.06±487.811 | <0.05 |
| Proteinuria (+) | ++~+++ | NA | NA |

sPE, severe pre-eclampsia; SBP, systolic blood pressure; DBP, diastolic blood pressure;
NA, not applicable. All data are presented as means±SD.

**Table S2.** Antibodies used for immunostaining and Western blot analysis.

| Antibody | Dilution | Source | Identifier |
| --- | --- | --- | --- |
| Rabbit polyclonal anti-Cytokeratin | 1/300 | DAKO | Z0622 |
| Rat polyclonal anti-CD31 | 1/300 | Biolegend | 102501 |
| Rabbit polyclonal anti-α-SMA | 1/200 | Proteintech | 55135-1-AP |
| Rabbit polyclonal anti-Axl | 1/1000 | Affinity | AF7793 |
| Rabbit polyclonal anti-p-STAT3(Ser727) | 1/1000 | Cell signaling technology | 49081 |
| Rabbit polyclonal anti- STAT3 | 1/1000 | Cell signaling technology | 4904 |
| Rabbit polyclonal anti-Corin | 1/500 | ABclonal technology | A10404 |
| Rabbit polyclonal anti-Gapdh | 1/1000 | Cell signaling technology | 5174 |

**Table S3.**
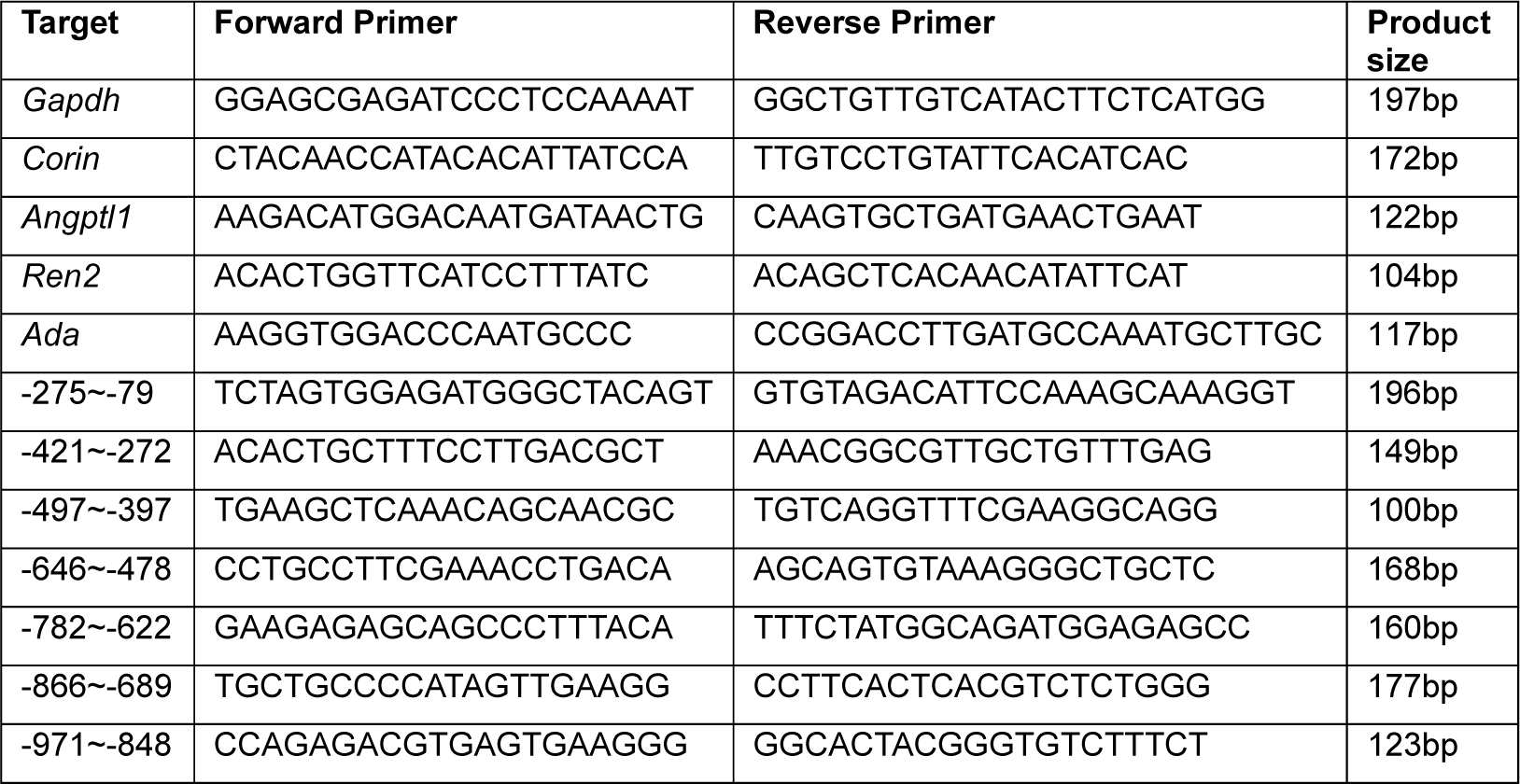
qPCR primers.

